# The biochemical basis of microRNA targeting efficacy

**DOI:** 10.1101/414763

**Authors:** Sean E. McGeary, Kathy S. Lin, Charlie Y. Shi, Namita Bisaria, David P. Bartel

## Abstract

MicroRNAs (miRNAs) act within Argonaute proteins to guide repression of mRNA targets. Although various approaches have provided insight into target recognition, the sparsity of miRNA–target affinity measurements has limited understanding and prediction of targeting efficacy. Here, we adapted RNA bind-n-seq to enable measurement of relative binding affinities between Argonaute–miRNA complexes and all ≤12-nucleotide sequences. This approach revealed noncanonical target sites unique to each miRNA, miRNA-specific differences in canonical target-site affinities, and a 100-fold impact of dinucleotides flanking each site. These data enabled construction of a biochemical model of miRNA-mediated repression, which was extended to all miRNA sequences using a convolutional neural network. This model substantially improved prediction of cellular repression, thereby providing a biochemical basis for quantitatively integrating miRNAs into gene-regulatory networks.

## INTRODUCTION

MicroRNAs (miRNAs) are ∼22-nt regulatory RNAs that derive from hairpin regions of precursor transcripts (*1*). Each miRNA associates with an Argonaute (AGO) protein to form a silencing complex, in which the miRNA pairs to sites within target transcripts and the AGO protein promotes destabilization and/or translational repression of bound target (*2*). miRNAs are grouped into families based on the sequence of their extended seed (nucleotides 2–8 of the miRNA), which is the region of the miRNA most important for target recognition (*3*). The 90 most broadly conserved miRNA families of mammals each have an average of >400 preferentially conserved targets, such that mRNAs from most human genes are conserved targets of at least one miRNA (*4*). Most of these 90 broadly conserved families are required for normal development or physiology, as shown by knockout studies in mice (*1*).

Deeper understanding of these numerous biological functions would be facilitated by a better understanding of miRNA targeting, with the ultimate goal of correctly predicting the effects of each miRNA on the output of each expressed gene. In principle, targeting efficacy should be a function of the affinity between target RNAs and AGO–miRNA complexes. However, binding affinities have been determined for only a few target sequences of only three miRNAs (*5*–*11*). These affinities show that the established parameters describing RNA-duplex stability in solution do not apply to RNA–RNA interactions in the context of AGO (*5, 6*). However, the sparsity of the biochemical data limits insight into how targeting might differ between different miRNAs and prevents construction of an informative biochemical model of targeting efficacy. Instead, the most informative of models of targeting efficacy rely on indirect inference through correlative approaches. These models focus on mRNAs with canonical 6–8-nt sites matching the miRNA seed region (Fig. 1A) and train on features known to correlate with targeting efficacy (including the type of site as well as various features of site context, mRNAs, and miRNAs), using datasets that monitor mRNA changes that occur after introducing a miRNA (*12*–*15*). Although the correlative model implemented in TargetScan7 performs well—as well as the best in vivo crosslinking approaches, it nonetheless explains only a small fraction of the mRNA changes observed upon introducing a miRNA (*r*^2^ = 0.14) (*13*). This low value indicates that prediction of targeting efficacy has room for improvement, even when accounting for the fact that experimental noise and secondary effects of inhibiting direct targets place a ceiling on the variability attributable to direct targeting. Therefore, we adapted RNA bind-n-seq (RBNS) (*16*) and a convolutional neural network (CNN) to the study of miRNA–target interactions, with the goal of obtaining the quantity and diversity of affinity measurements needed to better understand and predict miRNA targeting efficacy.

**Fig. 1.**
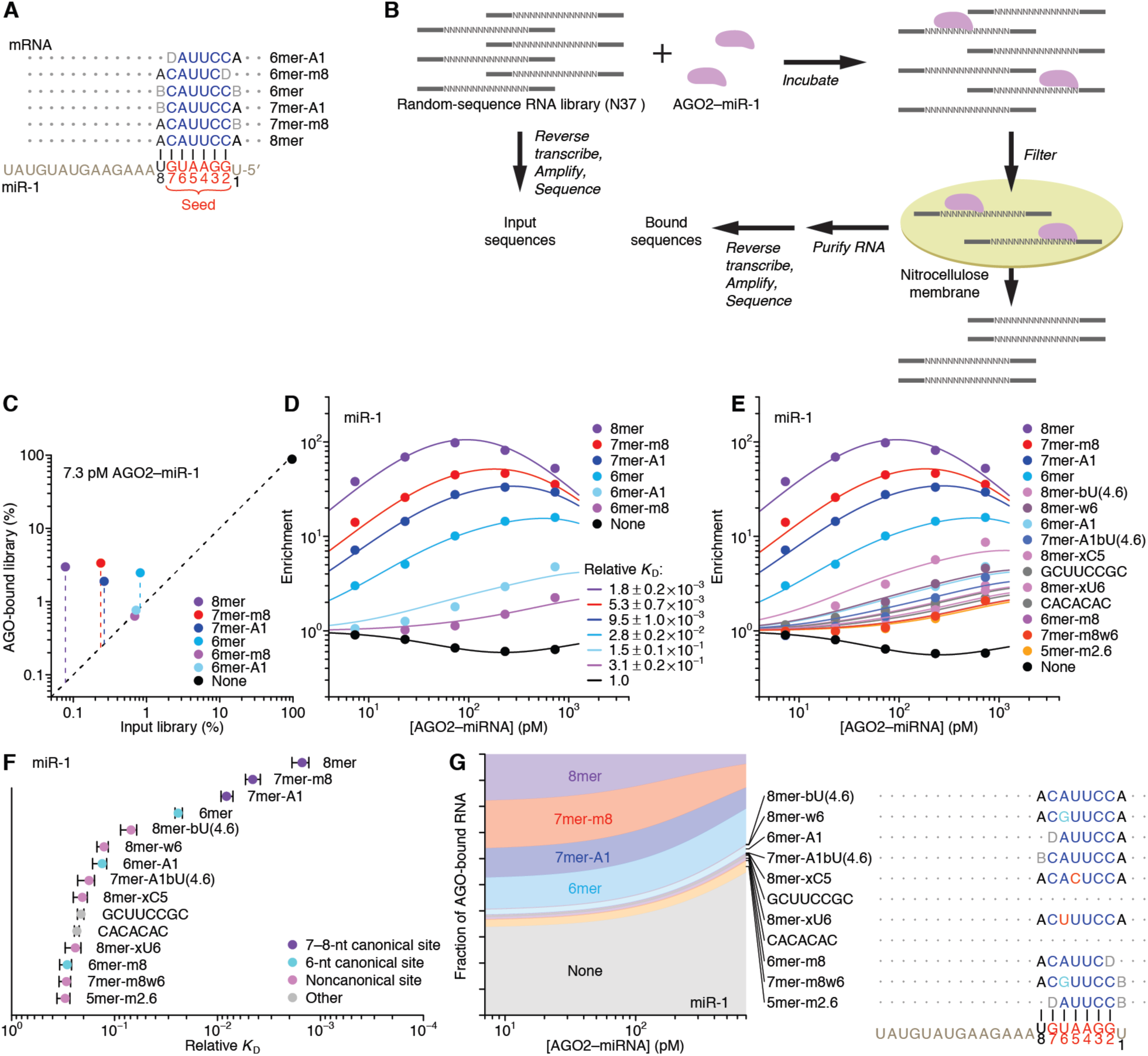
AGO-RBNS reveals binding affinities of canonical and novel miR-1 target sites. (**A**) Canonical sites of miR-1. These sites have contiguous pairing (blue) to the miRNA seed (red), and some include an additional match to miRNA nucleotide 8 or an A opposite miRNA nucleotide 1 (B, not A; D, not C). (**B**) AGO-RBNS. Purified AGO2–miR-1 is incubated with excess RNA library molecules that each have a central block of 37 random-sequence positions (N37). After reaching binding equilibrium, the reaction is applied to a nitrocellulose membrane and washed under vacuum to separate library molecules bound to AGO2–miR-1 from those that are unbound. Molecules retained on the filter are purified, reverse transcribed, amplified, and sequenced. These sequences are compared to those generated directly from the input RNA library. (**C**) Enrichment of reads containing canonical miR-1 sites in the 7.3 pM AGO2–miR-1 library. Shown is the abundance of reads containing the indicated site (key) in the bound library plotted as a function of the respective abundance in the input library. Dashed vertical lines depict the enrichment in the bound library; dashed diagonal line shows *y* = *x*. Reads containing multiple sites were assigned to the site with greatest enrichment. (**D**) AGO-RBNS profile of the canonical miR-1 sites. Plotted is the enrichment of reads with the indicated canonical site (key) observed at each of the five AGO2–miR-1 concentrations of the AGO-RBNS experiment, determined as in (C). Points show the observed values, and lines show the enrichment predicted from the mathematical model fit simultaneously to all of the data. Also shown for each site are *K*_D_ values obtained from fitting the model, listing the geometric mean ± the 95% confidence interval determined by resampling the read data, removing data for one AGO-miR-1 concentration and fitting the model to the remaining data, and repeating this procedure 200 times (40 times for each concentration omitted). (**E**) AGO-RBNS profile of the canonical and the newly identified noncanonical miR-1 sites (key). Sites are listed in the order of their *K*_D_ values and named and colored based on the most similar canonical site, indicating differences from this site with b (bulge), w (G–U wobble), or x (mismatch) followed by the nucleotide and its position. For example, the 8mer-bU(4.6) resembles a canonical 8mer site but has a bulged U at positions that would normally pair to miRNA nucleotides 4, 5, or 6. Otherwise, as in (D). (**F**) Relative *K*_D_ values for the canonical and the newly identified noncanonical miR-1 sites determined in (E). Sites are classified as either 7–8-nt canonical sites (purple), 6-nt canonical sites (cyan), noncanonical sites (pink), or a sequence motif with no clear complementarity to miR-1 (gray). The solid vertical line marks the reference *K*_D_ value of 1.0 assigned to reads lacking an annotated site. Error bars, 95% confidence interval on the geometric mean, as in (D). (**G**) The proportion of AGO2–miR-1 bound to each site type. Shown are proportions inferred by the mathematical model over a range of AGO2–miR-1 concentrations spanning the five experimental samples, plotted in the order of site affinity (top to bottom), using colors of (E). At the right is the pairing of each noncanonical site, diagrammed as in (A), indicating Watson–Crick pairing (blue), wobble pairing (cyan), mismatched pairing (red), bulged nucleotides (compressed rendering), and terminal non-complementarity (gray; B, not A; D, not C; H, not G; V, not U).

## RESULTS & DISCUSSION

### The site-affinity profile of miR-1

As previously implemented, RBNS provides qualitative relative binding measurements for an RNA-binding protein to a virtually exhaustive list of binding sites (*16, 17*). A purified RNA-binding protein is incubated with a large library of RNA molecules that each contain a central random-sequence region flanked by constant primer-binding regions. After reaching binding equilibrium, the protein is pulled down and any co-purifying RNA molecules are reverse transcribed, amplified, and sequenced. To extend RBNS to AGO–miRNA complexes (Fig. 1B), we purified human AGO2 loaded with miR-1 (*18*) (Fig. S1A) and set up five binding reactions, each with a different concentration of AGO2–miR-1 (range, 7.3–730 pM, logarithmically spaced) and a constant concentration of an RNA library with a 37-nt random-sequence region (100 nM). We also modified the protein-isolation step of the RBNS protocol, replacing protein pull-down with nitrocellulose filter binding, reasoning that the rapid wash step of filter binding would improve retention of low-affinity molecules that would otherwise be lost during the wash steps of a pull-down. This modified method was highly reproducible, with high correspondence observed between the 9-nt *k*-mer enrichments of two independent experiments using different preparations of both AGO2–miR-1 and RNA library (Fig. S1B; *r*^2^ = 0.86).

When analyzing our AGO-RBNS results, we first examined enrichment of the canonical miR-1 sites, comparing the frequency of these sites in RNA bound in the 7.3 pM AGO2–miR-1 sample with that of the input library. As expected from the site hierarchy observed in studies of site conservation and meta analyses of endogenous site efficacy (*3*), the 8mer site (perfect match to miR-1 nucleotides 2–8 followed by an A) was most enriched (38 fold), followed by the 7mer-m8 site (perfect match to miR-1 nucleotides 2–8, enrichment 14 fold), then the 7mer-A1 site (perfect match to miR-1 nucleotides 2–7 followed by an A, enrichment 7.2 fold), and the 6mer site (perfect match to miR-1 nucleotides 2–7, enrichment 3.0 fold) (Fig. 1A, C). Little if any enrichment was observed for either the 6mer-A1 site (perfect match to miR-1 nucleotides 2–6 followed by an A) or the 6mer-m8 site (perfect match to miR-1 nucleotides 3–8) at this lowest concentration of 7.3 pM AGO2–miR-1 (Fig. 1A, C), consistent with their weak signal in previous analyses of conservation and efficacy (*4, 13, 19*). Enrichment of sites was quite uniform across the span of the random-sequence region, which indicated minimal influence from either the primer-binding sequences or supplementary pairing to the 3′ region of the miRNA (Fig. S1D). Although sites with supplementary pairing can have enhanced efficacy and affinity (*3, 5, 20*), the minimal influence of supplementary pairing reflected the rarity of such sites in our library.

Analysis of enrichment of the six canonical sites across all five AGO2–miR-1 concentrations illustrated two hallmarks of this experimental platform (*16*). First, as the concentration increased from 7.3 pM to 73 pM, enrichment for each of the six site types increased (Fig. 1D), which was attributable to an increase in signal over a constant low background of library molecules isolated even in the absence of AGO2–miR-1. Second, as the AGO2–miR-1 concentration increased beyond 73 pM, 8mer enrichment decreased, and at the highest AGO2–miR-1 concentration, enrichment of the 7mer-m8 and 7mer-A1 site decreased (Fig. 1D). These waning enrichments indicated the onset of saturation for these high-affinity sites (*16*). These two features, driven by AGO–miRNA-independent background and partial saturation of the higher-affinity sites, respectively, caused differences in enrichment values for different site types to be highly dependent on the AGO2–miR-1 concentration; the lower AGO2–miR-1 concentrations provided greater discrimination between the higher-affinity site types, the higher AGO2–miR-1 concentrations provided greater discrimination between the lower-affinity site types, and no single concentration provided results that quantitatively reflected differences in relative binding affinities.

To account for background binding and ligand saturation, we developed a computational strategy that simultaneously incorporated information from all concentrations of an RBNS experiment to calculate relative *K*_D_ values. Underlying this strategy was an equilibrium-binding model that predicts the observed enrichment of each site type across the concentration series as a function of the *K*_D_ values for each miRNA site type (including the “no-site” type), as well as the stock concentration of purified AGO2–miR-1 and a constant amount of library recovered as background in all samples. Using this model, we performed maximum likelihood estimation (MLE) to fit the relative *K*_D_ values, which explained the observed data strikingly well (Fig. 1D). Moreover, these relative *K*_D_ values were robustly estimated, as indicated by comparing values obtained using results from only four of the five AGO2–miR-1 concentrations (*r*^2^ ≥ 0.994 for each of the ten pairwise comparisons, Fig. S1F, G). These quantitative binding affinities followed the same hierarchy as observed for site enrichment, but the differences in affinities were of greater magnitude (Fig. 1D, Fig. S1C).

Up to this point, our analysis was informed by the wealth of previous computational and experimental data showing the importance of a perfect 6–8-nt match to the seed region (*3*). However, the ability to calculate the relative *K*_D_ of any *k*-mer of length ≤12 nt (the 12-nt limit imposed by the sparsity of reads with longer *k*-mers) provided the opportunity for a de novo search for sites, without requiring complementarity to the miRNA sequence and without bias from any previous knowledge. In this search, we 1) calculated the enrichment of all 10-nt *k*-mers in the bound RNA in the 730 pM AGO2–miR-1 sample, which was the sample with the most sensitivity for detecting low-affinity sites, 2) determined the extent of complementarity of the top 10 *k*-mers to the miR-1 sequence, 3) assigned a site most consistent with the observed *k*-mers, and 4) removed all reads containing this newly identified site from both the bound and input libraries. These four steps were iterated until no 10-nt *k*-mers remained that were enriched ≥10-fold, thereby generating 15 sites for AGO2–miR-1. We then applied our MLE procedure to calculate relative *K*_D_ values for this expanded list of sites (Fig. 1E, F).

This unbiased approach demonstrated that the 8mer, 7mer-m8, 7mer-A1, and 6mer sites to miR-1 were the highest-affinity site types of lengths ≤10 nt, and also identified 9 novel site types with binding affinities resembling those of the 6mer-m8 and the 6mer-A1 (Fig. 1F). Comparison of these sites to the sequence of miR-1 revealed that miR-1 can tolerate either a wobble G at position 6 or a bulged U somewhere between positions 4 and 6 and achieve affinity at least 7–11 fold above the remaining no-site reads. We also identified two motifs with enrichment more difficult to attribute solely to miRNA pairing. These were the GCUUCCGC motif, which had contiguous complementarity to positions 2–5 of miR-1 flanked by noncomplementary GC dinucleotides on both sides, and the CACACAC motif, which had contiguous complementarity that did not extend beyond the UGU at positions 6–8 and 18–20 of miR-1. Nonetheless, among the 1,398,100 possible motifs ≤10 nt, these were the only two that passed our cutoffs and were difficult to attribute to miRNA pairing.

Our analytical approach and its underlying biochemical model also allowed us to infer the proportion of AGO2–miR-1 bound to each site (Fig. 1G). The 8mer site occupied 3.6–18% of the silencing complex over the concentration course, whereas the 7mer-m8, by virtue of its greater abundance occupied a somewhat greater fraction of the complex. In aggregate, the marginal sites, including the 6mer-A1, 6mer-m8, and nine noncanonical sites, occupied 7–11% of the AGO2–miR-1 complex. Moreover, because of their very high abundance, library molecules with no identified site occupied 33–55% of the complex (Fig. 1G). These results support the inference that the summed contributions of background binding and low-affinity sites to intracellular AGO occupancy is of the same order of magnitude as that of canonical sites, suggesting that an individual AGO–miRNA complex spends about half its time associated with a vast repertoire of background and low-affinity sites (*21, 22*).

Our results confirmed that AGO2–miR-1 binds the 8mer, 7mer-m8, 7mer-A1, and 6mer sites most effectively and revealed the relative binding affinities and occupancies of these sites. In addition, our results uncovered weak yet specific affinity to the 6mer-A1 and 6mer-m8 sites plus nine noncanonical sites. Although alternative binding sites for miRNAs have been proposed based on high-throughput in vivo crosslinking studies (*23*–*27*), our approach provided quantification of the relative strength of these sites without the confounding effects of differential crosslinking efficiencies, potentially enabling their incorporation into a quantitative framework of miRNA targeting.

### Distinct canonical and noncanonical binding of different miRNAs

We extended our analysis to five additional miRNAs, including let-7a, miR-7, miR-124, and miR-155 of mammals (Fig. 2, Fig. S2), chosen for their sequence conservation as well as the availability of data examining their regulatory activities, intracellular binding sites, or in vitro binding affinities (*1, 5, 6, 23, 24*), and lsy-6 of nematodes, which is thought to bind unusually weakly to its canonical sites (*28*). In the case of let-7a, previous biochemical analyses have determined the *K*_D_ values of a few sites (*5, 6*), and our values agreed well, which further validated our high-throughput approach (Fig. S1H).

**Fig. 2.**
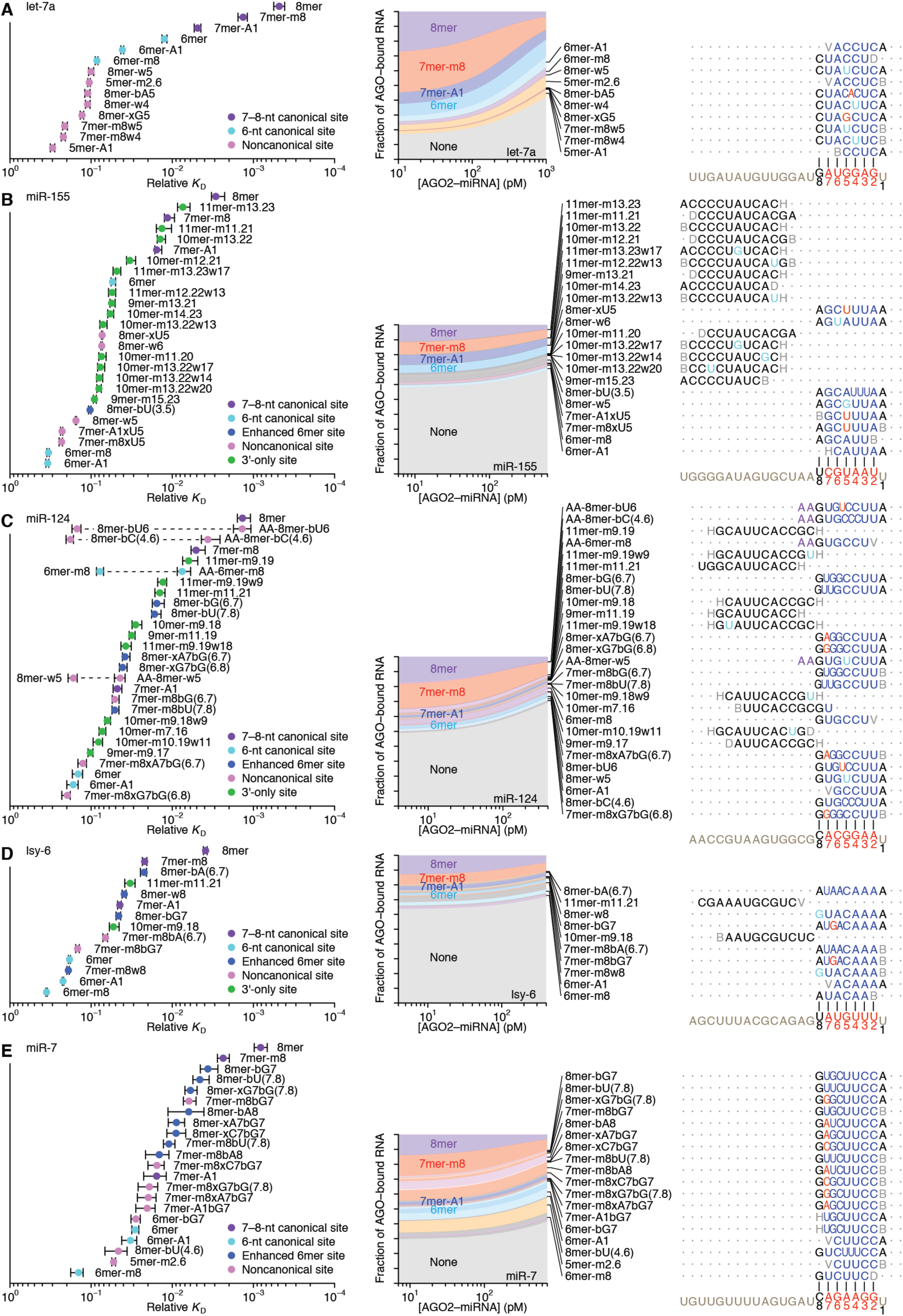
Distinct canonical and noncanonical binding of different miRNAs. (**A**–**E**) Relative *K*_D_ values and proportional occupancy of established and newly identified sites of let-7a (A), miR-155 (B), miR-124 (C), lsy-6 (D), and miR-7 (E). To conserve space, sites with no clear complementarity to the miRNA are omitted, as are most miR-124 sites extended with 5′-AA. A complete list of sites is provided in Fig. S2. The 5′-AA-extended sites that are shown are the four that were extensions of sites that were also identified without extensions; for these four, both the smaller site and its 5′-AA-extended version are shown on the same line, connected with a dashed line (C). Relative *K*_D_ values are plotted as in Fig. 1F but in some cases with additional categories, either for 3′-only sites (green) (B–D) or for 6-nt canonical sites enhanced by either additional wobble-pairing or additional Watson–Crick complementarity separated by a bugled nucleotide (blue) (B–E). The proportion of AGO2–miRNA bound to each site type (including those listed only in Fig. S2) is estimated and shown as in Fig. 1G.

The site-affinity profile of let-7a resembled that of miR-1, except the 6mer-m8 and 6mer-A1 sites for let-7a had greater binding affinity than essentially all of the noncanonical sites (Fig. 2A, Fig. S2A). As with miR-1, the noncanonical sites each paired to the seed region but did so imperfectly, typically with a single wobble, single mismatch, or single-nucleotide bulge, but these imperfections differed from those observed for miR-1 (Figs. 1F, 2A). The let-7a analysis also identified a site that, as with the miR-1 CGUUCCGC and CACACAC sites, could not be explained by pairing to the miRNA (Fig. S2A). These rare sites that lacked substantial pairing to the miRNA always differed for different miRNAs, which ruled out binding to a common contaminant in our AGO2–miRNA preparations (Fig. 1F, Fig. S2).

The site-affinity profiles of miR-124, miR-155, lsy-6, and miR-7 resembled those of miR-1 and let-7a. They all included the six canonical sites and noncanonical sites with extensive yet imperfect pairing to the miRNA seeds, with these imperfections tending to occur at different positions for different miRNAs, with different mismatched-or bulged-nucleotide identities (Fig. 2B–E, Fig. S2). In contrast to the noncanonical sites of miR-1 and let-7a, more of the noncanonical sites of the other four miRNAs had affinities interspersed with those of the top four canonical sites. Moreover, the profiles for miR-155, miR-124, and lsy-6 also included sites with extended (9–11-nt) complementarity to the miRNA 3′ region. These sites had estimated *K*_D_ values that were derived from reads with little more than chance complementarity to the miRNA seed, and they had uniform enrichment across the length of the random-sequence region (Fig. S1E), which indicated that these sites represented an alternative binding mode dominated by extensive pairing to the 3′ region without involvement of the seed region (Fig. 2B–D, Fig. S2B– D). We named them “3′-only sites”.

In some respects the 3′-only sites resembled noncanonical sites known as centered sites, which are reported to function in mammalian cells (*29*). Like 3′-only sites, centered sites have extensive perfect pairing to the miRNA, but for centered sites this pairing begins at miRNA positions 3 or 4 and extends 11–12-nt through the center of the miRNA (*29*). Our unbiased search for sites did not identify centered sites for any of the six miRNAs. We therefore directly queried the region of each miRNA to which extensive noncanonical pairing was favored, determining the affinity of sequences with 11-nt segments of perfect complementarity to the miRNA sequence, scanning from miRNA position 3 to the 3′ end of the miRNA (Fig. 3A). For miR-155, miR-124, and lsy-6, sequences with 11-nt sites that paired to the miRNA 3′ region bound with greater affinity than did those with a canonical 6mer site, whereas for let-7a and miR-1, and miR-7, none of the 11-nt sites conferred stronger binding than did the 6mer. Moreover, for all six miRNAs, the 11-nt sites that satisfied the criteria for annotation as centered sites conferred binding ≤2-fold stronger than that of the 6mer-m8 site, which also starts at position 3 but extends only 6 nt. These results called into question the function of centered sites, although we cannot rule out the possibility that centered sites are recognized by some miRNAs and not others. Indeed, the newly identified 3′-only sites functioned for only miR-155, miR-124, and lsy-6, and even among these, the optimal region of pairing differed, occurring at positions 13–23, 9–19, and 8–18, respectively (Fig. 3A).

**Fig. 3.**
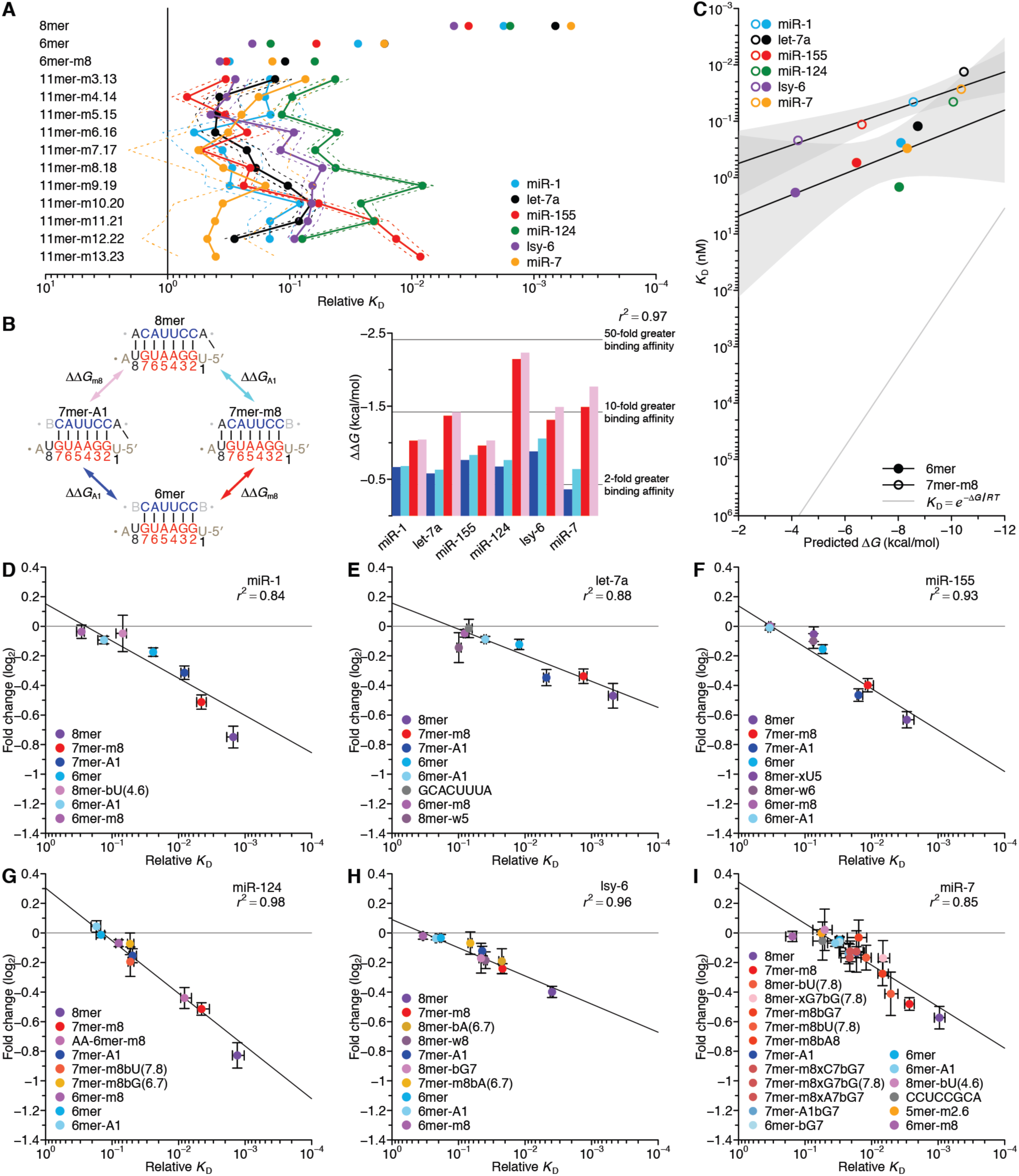
Additional analyses of binding affinities and the correspondence between binding affinity and repression efficacy. (**A**) Diverse functionality and position dependence of 11-nt 3′-only sites. Relative *K*_D_ values for each potential 11-nt 3′-only site are plotted for the indicated miRNAs (key). For reference, values for the 8mer, 6mer, and 6mer-m8 sites are also plotted. The solid vertical line marks the reference *K*_D_ value of 1.0, as in Fig. 1F. The solid and dashed lines indicate geometric mean and 95% confidence interval, respectively, determined as in Fig. 1D. (**B**) The independent contributions of the A1 and m8 features. At the left a double mutant cycle depicts the affinity differences observed among the four top canonical sites for miR-1, as imparted by the independent contributions of the A1 and m8 features and their potential interaction. At the right the apparent binding contributions of the A1 (ΔΔ*G*_A1_, blue and cyan) or m8 (ΔΔ*G*_m8_, red and pink) features are plotted, determined from the ratio of relative *K*_D_ values of either the 7mer-A1 and the 6mer (blue), the 8mer and the 7mer-m8 (cyan), the 7mer-m8 and the 6mer (red), or the 8mer and the 7mer-A1 (pink), for the indicated AGO2–miRNA complexes. The *r*^2^ reports on the degree of ΔΔ*G* similarity for both the m8 and A1 features using either of the relevant site-type pairs across all six complexes. (**C**) The relationship between the observed relative *K*_D_ values and predicted pairing stability of the 6mer (filled circles) and 7mer-m8 (open circles) sites of the indicated AGO–miRNA complex (key), under the assumption that the no-site *K*_D_ (set to 10 nM for modeling and *K*_D_ estimation) was consistent across AGO–miRNA complexes. The two black lines are the best fit of the relationship observed for each of the site types (gray regions, 95% confidence interval, for linear regression predicting log-transformed *K*_D_ values). The gray line shows the expected relationship with the predicted stabilities given by *K*_D_ = *e*^-Δ*G*/*RT*^. **(D–I**) The relationship between repression efficacy and relative *K*_D_ values for the indicated sites of miR-1 (D), let-7a (E), miR-155 (F), miR-124 (G), lsy-6 (H), and miR-7 (I). To include information from mRNAs with multiple sites, multiple linear regression was applied to determine the log fold-change attributable to each site type (error bars, 95% confidence interval). The relative *K*_D_ values are those of Figs. 1 and 2 (error bars, 95% confidence interval). Lines show the best fit to the data as determined by least-squares regression with the log-transformed *K*_D_ values, weighting residuals using the 95% confidence intervals of the log fold-change estimates. The *r*^2^ values were calculated using similarly weighted Pearson correlations.

When evaluating other types of noncanonical site proposed to confer widespread repression in mammalian cells (*19, 23*), we found that all but two bound with affinities difficult to distinguish from background. One of these two was the 5-nt site matching miRNA positions 2–6 (5mer-m2.6) (*19*), which was bound by miR-1, let-7a, and miR-7 but not by the other three miRNAs (Fig. S3). The other was the pivot site (*23*), which was bound by miR-124 (e.g., 8mer-bG(6.7); Fig. 2C) and lsy-6 (e.g., 8mer-bA(6.7); Fig. 2D) but not by the other four miRNAs (Fig. S4). Thus, these two previously identified noncanonical site types resembled the newly identified noncanonical sites with extensive yet imperfect pairing to the seed region, in that they function for only limited number of miRNAs.

In addition to the differences in noncanonical site types observed for each miRNA, we also observed striking miRNA-specific differences in the relative affinities of the canonical site types. For example, for miR-155, the affinity of the 7mer-A1 nearly matched that of the 7mer-m8, whereas for miR-124, the affinity of the 7mer-A1 was >11-fold lower than that of the 7mer-m8. These results implied that the relative contributions of the A at target position 1 and the match at target position 8 can substantially differ for different miRNAs. Although prior studies show that AGO proteins remodel the thermodynamic properties of their loaded RNA guides (*5, 6*), our results show that the sequence of the guide strongly influences the nature of this remodeling, leading to differences in relative affinities across canonical site types and a distinct repertoire of noncanonical site types for each miRNA.

### The energetics of canonical binding

With the relative *K*_D_ values for the canonical binding sites of six miRNAs in hand, we examined the energetic relationship between the A at target position 1 (A1) and the match at miRNA position 8 (m8), within the framework analogous to a double-mutant cycle (Fig. 3B, left). The apparent binding-energy contributions of the m8 and A1 (ΔΔ*G*_m8_ and ΔΔ*G*_A1_, respectively) were largely independent, as inferred from the relative *K*_D_ values of the four site types. That is, for each miRNA, the ΔΔ*G*_m8_ inferred in presence of the A1 (using the ratio of the 8mer and 7mer-A1 *K*_D_ values) resembled that inferred in the absence of the A1 (using the ratio of the 7mer-m8 and 6mer *K*_D_ values), and vice versa (Fig. 3B).

The relative *K*_D_ values for canonical sites of six miRNAs provided the opportunity to examine the relationship between the predicted free energy of site pairing and measured site affinities. We focused on the 6mer and 7mer-m8 sites, because they lack the A1, which does not pair to the miRNA (Fig. 1A) (*8, 30*). Consistent with the importance of base pairing for site recognition and the known relationship between predicted seed-pairing stability and repression efficacy (*28*), affinity increased with increased predicted pairing stability, although this increase was statistically significant for only the 7mer-m8 site type (Fig. 3C, *p =* 0.12 and 0.006, for the 6mer and 7mer-m8 sites, respectively). However, for both site types, the slope of the relationship was significantly less than expected from *K*_D_ = *e*^-Δ*G*/*RT*^ (*p =* 0.005 and 5 × 10^-5^, respectively). When considered together with previous analysis of a miRNA with enhanced seed pairing stability, these results indicated that in remodeling the thermodynamic properties of the loaded miRNAs, AGO not only enhances the affinity of seed-matched interactions but also dampens the intrinsic differences in seed-pairing stabilities that would otherwise impose much greater inequities between the targeting efficacies of different miRNAs (*6*). Thus, although lsy-6, which has unusually poor predicted seed-pairing stability (*28*), did indeed have the weakest site-binding affinity of the six miRNAs, the difference between its binding affinity and that of the other miRNAs was less than might have been expected.

### Correspondence between affinity measured by AGO-RBNS and repression observed in the cell

To evaluate the relevance of our in vitro binding results to intracellular miRNA-mediated repression, we examined the relationship between the relative *K*_D_ measurements and the repression of endogenous mRNAs after miRNA transfection into HeLa cells. When examining intracellular repression attributable to 3′-UTR sites to the transfected miRNA, we observed a striking relationship between AGO-RBNS–determined *K*_D_ values and mRNA fold-changes (Fig. 3D–I, *r*^2^ = 0.84–0.98). For instance, the different relative affinities of the 7mer-A1 and 7mer-m8 sites, most extremely observed for sites of miR-155 and miR-124, was nearly perfectly mirrored by the relative efficacy of these sites in mediating repression in the cell (Fig. 3F, G). A similar correspondence between relative *K*_D_ values and repression was observed for the noncanonical sites that had both sufficient affinity and sufficient representation in the HeLa transcriptome to be evaluated using this analysis (Fig. 3D–I). These included the pivot sites for miR-124 and lsy-6, the AA-6mer-m8 site for miR-124, and the bulge-G7-containing sites for lsy-6 and miR-7 (Fig. 3G–I). The consistent relationship between in vitro binding affinity and intracellular repression supported a model in which repression is a function of miRNA occupancy, as dictated by site affinity, and thus miRNA-and site-specific differences in binding affinities explain substantial differences in repression.

### The strong influence of flanking dinucleotide sequences

AU-rich nucleotide composition immediately flanking miRNA sites has long been associated with increased site conservation and efficacy in cells (*12, 30, 31*), but the mechanistic basis of this phenomenon had not been investigated, presumably because of the sparsity of affinity measurements. The AGO-RBNS data provided the means to overcome this limitation. We first separated the miR-1 8mer site into 256 different 12-nt sites, based on the dinucleotide sequences immediately flanking each side of the 8mer, and determined relative *K*_D_ values for each (Fig. 4A). This analysis revealed a 100-fold range in values, depending on the identities of the flanking dinucleotides (1,840-and 17-fold from background for AU-8mer-UA and GG-8mer-GG, respectively), with binding affinity strongly tracking the AU content of the flanking dinucleotides. Extending this analysis across all miR-1 site types (Fig. 4B), as well as to sites to the other five miRNAs (Fig. S5A–E), yielded similar results. The effect of flanking-dinucleotide context was of such magnitude that it often exceeded the affinity differences observed between miRNA-site types. Indeed, for each miRNA, at least one 6-nt canonical site in its most favorable context had greater affinity than that of the 8mer site in its least favorable context (Fig. 4B, Fig. S5A–E).

**Fig. 4.**
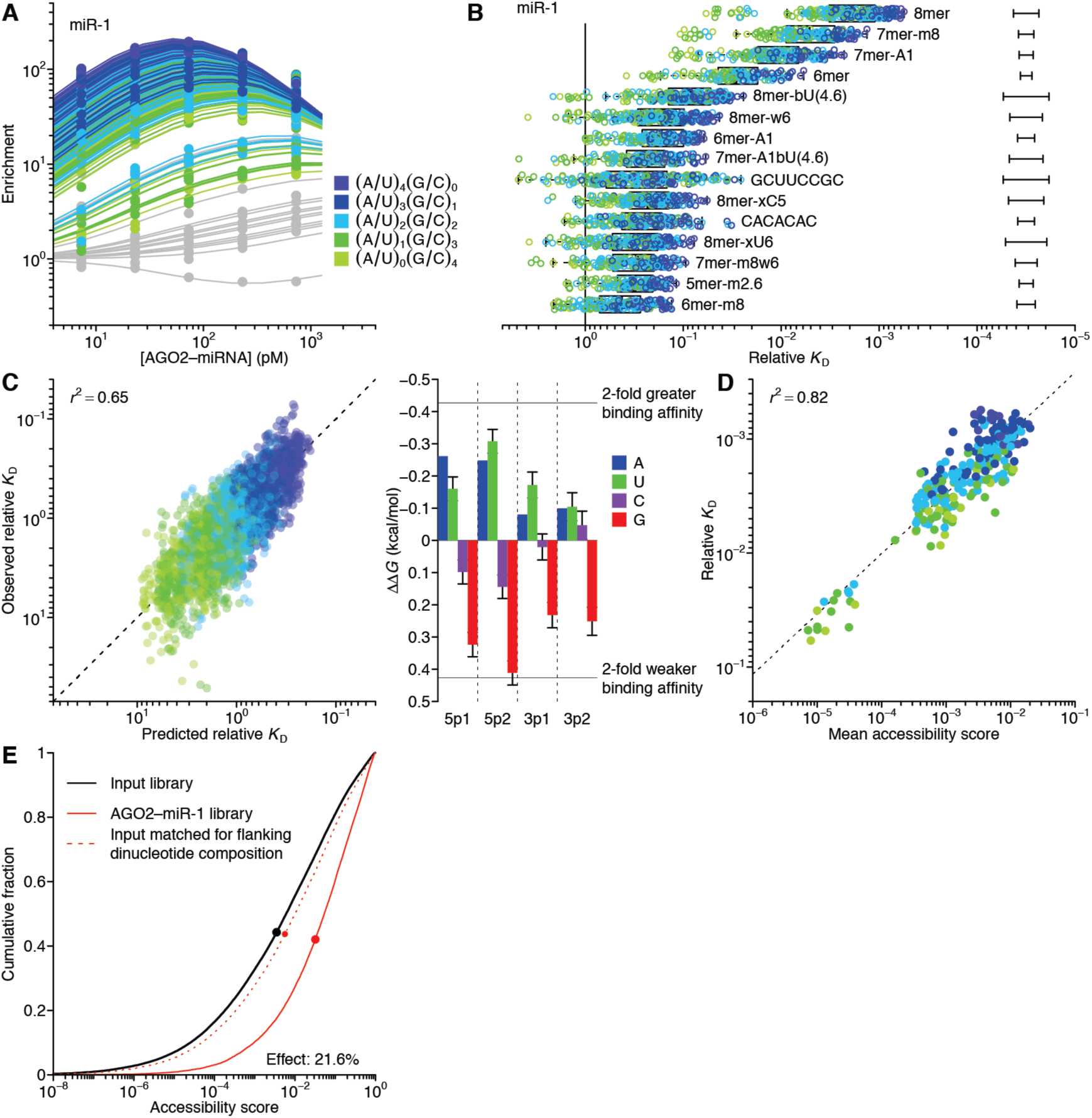
The influence of flanking dinucleotide sequence context. (**A**) AGO-RBNS profile of miR-1 sites, showing results for the 8mer separated into 256 different 12-nt sites based on the identities of the two dinucleotides immediately flanking the 8mer. For each 12-nt site, the points and line are colored based on the AU content of the flanking dinucleotides (key). For context, results of Fig. 1E are re-plotted in gray. Otherwise as in Fig. 1E. (**B**) Relative *K*_D_ values for the each miR-1 site identified in Fig. 1F separated into 144–256 sites as in (A) based on the identities of the flanking dinucleotides. The points are colored as in (A). Error bars, median 95% confidence interval across all *K*_D_ values. Otherwise, as in Fig. 1F. (**C**) Consistency of flanking-dinucleotide effect across miRNA and site type. At the left is a comparison of observed relative *K*_D_ values and results of a mathematical model that used multiple linear regression to predict the influence of flanking dinucleotides on log-transformed *K*_D_. Plotted are results for all flanking dinucleotide contexts of all six canonical site types, for all six miRNAs, normalized to the average affinity of each canonical site. Predictions of the model are those observed in a six-fold cross validation, training on the results for five miRNAs and reporting the predictions for the held-out miRNA. The *r*^2^ quantifies the agreement between the log-transformed predicted and actual values. At the right the model coefficients (multiplied by -*RT*, where *T* = 310.15 K) corresponding to each of the four nucleotides of the 5′ (5p) and 3′ (3p) dinucleotides in the 5′-to-3′ direction are plotted (error bars, 95% confidence interval). (**D**) Relationship between the mean structural-accessibility score and the relative *K*_D_ for the 256 12-nt sites containing the miR-1 8mer flanked by each of the dinucleotide combinations. Points are colored as in (A). Linear regression (dashed line) and calculation of *r*^2^ were performed using log-transformed values. For an analysis of the relationship between 8mer flanking dinucleotide *K*_D_ and structural accessibility over a range of window lengths and positions relative to the 8mer site, see Fig. S5G. (**E**) The influence of site accessibility after accounting for nucleotide sequence composition. Plotted are cumulative distributions of structural-accessibility scores of the 8mer sites of the input library (solid black line), 8mer sites of the bound library from the 7.3 nM sample (solid red line), and 8mer sites of 8mer-containing reads from the input library sampled to match the flanking dinucleotide composition of the 8mer sites in bound the 7.3 nM sample (dashed red line). The geometric mean of each distribution is indicated (points). The geometric mean of the resampled distribution spanned 21.6% of the logarithmic difference in geometric means observed between the two experimental distributions.

To identify general features of the flanking-dinucleotide effect across miRNA sequences and site types, we trained a multiple linear-regression model on the complete set of flanking-dinucleotide *K*_D_ values corresponding to all six canonical site types of each miRNA, fitting the effects at each of the four positions within the two flanking dinucleotides. The output of the model agreed well with the observed *K*_D_ values (Fig. 4C, left, *r*^2^ = 0.65), which indicated that the effects of the flanking dinucleotides were largely consistent between miRNAs and between site types of each miRNA. A and U nucleotides each enhanced affinity, whereas G nucleotides reduced affinity, and C nucleotides were intermediate or neutral (Fig. 4C, right). Moreover, the identity of the 5′ flanking dinucleotide, which must come into close proximity with the central RNA-binding channel of AGO (*7*), contributed ∼2-fold more to binding affinity than did the 3′ flanking sequence (Fig. 4C, right).

One explanation for this hierarchy of flanking nucleotide contributions, with A ≈ U > C > G, is that it inversely reflected the propensity of these nucleotides to stabilize RNA secondary structure that could occlude binding of the silencing complex. To investigate this potential role for structural accessibility in influencing binding, we compared the predicted structural accessibility of 8mer sites in the input and bound libraries of the AGO2–miR-1 experiment, using a score for predicted site accessibility previously optimized on data examining miRNA-mediated repression (*13, 32*). This score is based on the predicted probability that the 14-nt segment at target positions 1–14 is unpaired. We found that predicted accessibilities of sites in the bound libraries were substantially greater than those for sites in the input library, and the difference was greatest for the samples with the lower AGO2–miR-1 concentrations (Fig. S5F), as expected if the accessibility score was predictive of site accessibility and if the most accessible sites were the most preferentially bound.

To build on these results, we examined the relationship between predicted structural accessibility and binding affinity for each of the 256 flanking dinucleotide possibilities. For each input read with a miR-1 8mer site, the accessibility score of that site was calculated. The sites were then differentiated based on their flanking dinucleotides into 256 12-nt sites, and the geometric mean of the site-accessibility scores of each of these extended sites was compared with the AGO-RBNS–derived relative *K*_D_ value (Fig. 4D, Fig. S5G). A striking correlation was observed (*r*^2^ = 0.82, *p* < 10^-15^), with all 16 sites containing a 5′-flanking GG dinucleotide having both unusually poor affinities and unusually low accessibility scores.

Although we cannot rule out the possibility that the flanking dinucleotide preferences are caused by direct contacts to AGO with sequence preferences that happen to correlate strongly with those predicted structural accessibility, the high correspondence of predicted site accessibility and relative *K*_D_—one being the averaged result of a computational algorithm applied to reads from the input library, the other being a biochemical constant derived from RBNS analyses—strongly implied that site accessibility was the primary cause of the different binding affinities associated with flanking-dinucleotide context. Supporting this interpretation, we found that when the 8mer-containing reads of the input library were sampled to match the flanking dinucleotide distribution of the 8mer-containing reads in the 7.3 pM AGO2–miR-1 library, flanking dinucleotide identities explained only a minor fraction of the enrichment of structurally accessible reads observed in the bound libraries (Fig. 4E). Extending the analysis to data from the other four AGO2–miR-1 concentrations yielded consistent results, with the results from matched sampling of flanking dinucleotides never explaining >25% of the increased mean accessibility score (Fig. S5H). By contrast, sampling 8mer-containing reads from the input to match the site-accessibility scores of the bound reads yielded flanking dinucleotide frequencies that corresponded to those of the bound library (*r*^2^ = 0.79) (Fig. S5H). Taken together, our results demonstrate that local sequence context has a large influence on miRNA–target binding affinity, and indicate that this influence results predominantly from the differential propensities of flanking sequences to form structures that occlude site accessibility.

### A biochemical model predictive of miRNA-mediated repression

When predicted seed-pairing stability is included as a feature in target-prediction algorithms, it makes only a minor contribution to prediction accuracy (*13, 14*), suggesting that either target-site affinity is poorly represented by the predicted free energy of pairing or it is not the primary driver of repression efficacy. We found that predicted seed-pairing stability only poorly reflected actual affinity (Fig. 3C) and that measured affinities strongly corresponded to the repression observed in cells (Fig. 3D–I), which indicated that binding affinity is in fact a major determinant of targeting efficacy. With this in mind, we set out to build a biochemical framework of targeting efficacy. Previous efforts have used biochemical principles when modeling aspects of miRNA biology, including competition between endogenous target sites (*22, 33, 34*) and the influence of miRNAs on reporter gene–expression noise (*35*), but these efforts were severely limited by the sparsity of the data. Our ability to measure the relative binding affinity of a miRNA to any 12-nt sequence enabled modeling of the quantitative effects the six miRNAs on each cellular mRNA.

We first re-analyzed all six AGO-RBNS experiments to calculate, for each miRNA, the relative *K*_D_ values for all 262,144 12-nt *k*-mers that contained at least four contiguous nucleotides of the canonical 8mer site (Fig. 5A). These potential binding sites included the canonical sites and most of the noncanonical sites that we had identified, each within a diversity of flanking sequence contexts (Figs. 1F, 2). For each mRNA *m* and transfected miRNA *t*, the steady-state occupancy *N*_*m,t*_ (i.e., average number of AGO–miRNA complexes loaded with miRNA *t* bound to mRNA *m*) was predicted as a function of the *K*_D_ values of the potential binding sites contained within the mRNA 3′ UTR, as well as the concentration of the unbound AGO–miRNA_*t*_ complex *a*_*t*_, which was fit as a single value for each transfected miRNA (Fig. 5B). This occupancy value enabled prediction of a biochemically informed expectation of repression, assuming that added effect of the miRNA on the basal decay rate scaled with the basal rate and *N*_*m,t*_ (Fig. 5B, equation 3).

**Fig. 5.**
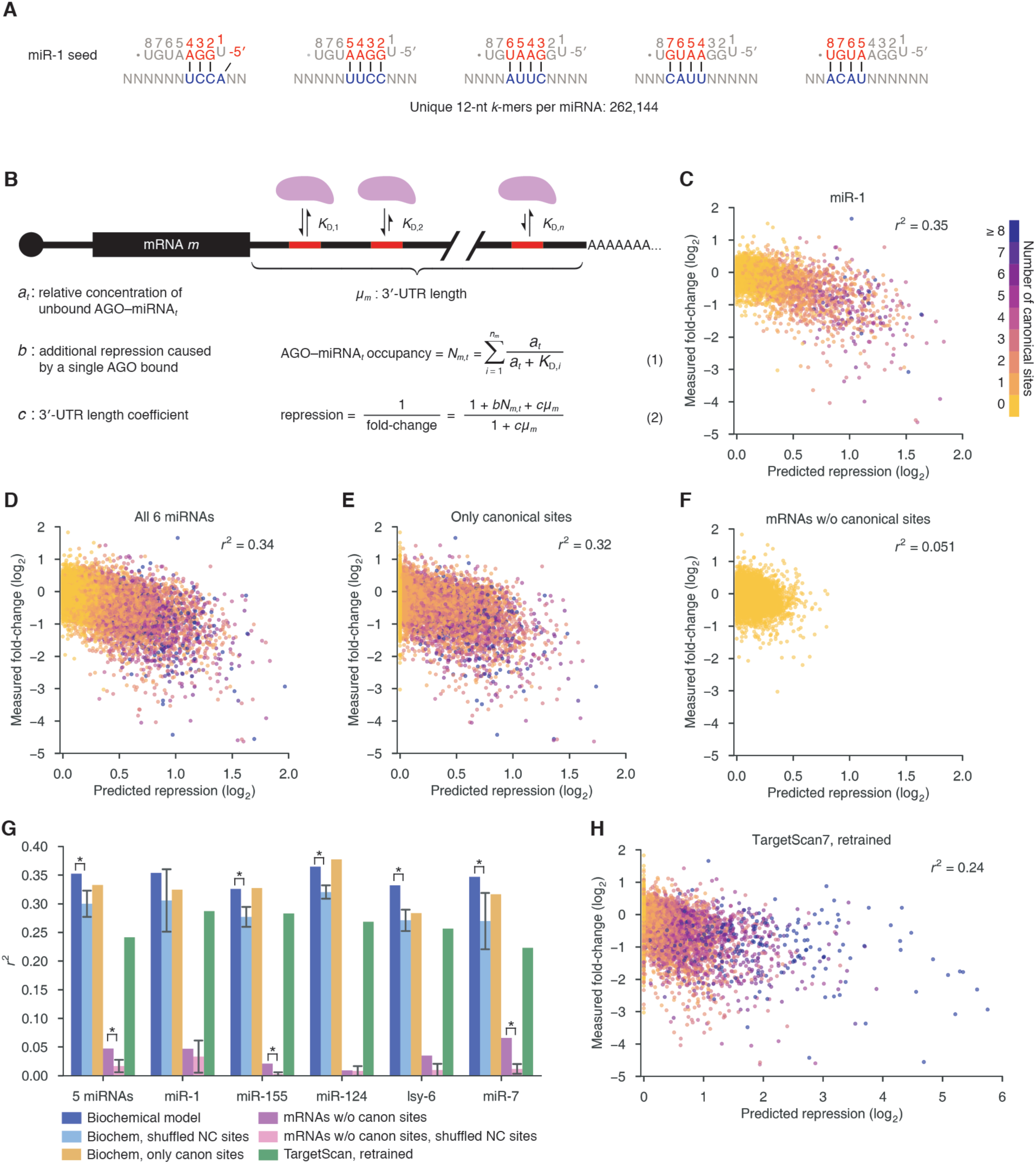
AGO-RBNS *K*_D_ values enable a predictive model of miRNA-mediated repression in cells. (**A**) The 262,144 12-nt *k*-mers with at least four contiguous matches to the extended seed region of miR-1, for which relative *K*_D_ values were determined. Relative *K*_D_ values were similarly determined for the analogous *k*-mers of the other five miRNAs. (**B**) Biochemical model for estimating the repression of an mRNA by a transfected miRNA using the relative *K*_D_ values of the 12-nt *k*-mers in the 3′ UTR of the mRNA. (**C**) Typical performance of the biochemical model. Results are shown for miR-1, chosen because among the six miRNAs tested, it had a median *r*^2^ value (Fig. S6A). Plotted is the relationship between mRNA changes observed after transfecting a miRNA and those predicted by the model. Each point represents the mRNA from one gene after transfection of miR-1 and is colored according to the number of canonical sites in the mRNA 3′ UTR (key). The Pearson’s *r*^2^ between measured and predicted values is reported in the upper right. (**D**) Performance of the biochemical model as evaluated using the combined results of all six miRNAs. Otherwise, as in (C). (**E**) Performance of the biochemical model when provided the relative *K*_D_ values of only the 12-nt *k*-mers that contain one of the six canonical site-types. Otherwise, as in (D). (**F**) Performance of the biochemical model when only considering mRNAs without one of the six canonical sites. Likelihood ratio test comparing this model to an intercept-only model with 8 degrees of freedom (for the 8 parameters fit by the biochemical model) yielded *p* < 10^-15^. Otherwise, as in (D). (**G**) The contributions of cognate noncanonical sites to performance of the biochemical model and comparison of its performance to that of TargetScan. Results are plotted for each of the miRNAs indicated and for all five miRNAs combined. To examine the contribution of noncanonical sites, datasets were shuffled by substituting the noncanonical sites (and corresponding relative *K*_D_ values) of each miRNA with those of a different miRNA. Shuffled datasets were generated for all 265 possible permutations, and for each permutation, the biochemical model was refit and evaluated. Plotted next to the performance (*r*^2^) observed for the indicated miRNA(s) (as in C or D, blue) is the average performance observed with shuffled noncanonical sites (light blue; error bars, standard deviation; asterisk, none of 265 model fits using shuffled noncanonical sites out-performed the model using the cognate noncanonical sites), and plotted next to that is the performance using only canonical sites (as in E, gold). An analogous analysis focused on only mRNAs without canonical sites, using either the cognate noncanonical sites (as in F, violet) or the average performance with shuffled noncanonical sites (pink, error bars, standard deviation; asterisk, none of 265 model fits using shuffled noncanonical sites out-performed the model using the cognate noncanonical sites). Also plotted is the test performance of the TargetScan model retrained on the 16 transfection datasets (green). (**H**) Test performance of the TargetScan model retrained on the 16 transfection datasets, evaluated using the combined results for the five miRNAs of (G). The *x*-axis was truncated at 6, which excluded two outlier points; otherwise, as in (C).

The calculation of predicted repression required an estimate of how much a single bound RISC complex affected the mRNA decay rate (Fig. 5B, *b*), which was fit as a global value for all six transfection experiments. Additionally, to account for the observation that longer 3′ UTRs tend to be less sensitive to miRNA-mediated repression (*13*), our model included a dampening term that scaled with the length of the mRNA 3′ UTR.

Our model was fit against repression observed in HeLa cells transfected with one of the six miRNAs. For five of the six miRNAs, a strong correspondence was observed when comparing mRNA changes measured upon miRNA transfection to those predicted by the model (Fig. 6C and Fig. S6A, *r*^2^ = 0.33–0.36). Performance was somewhat less for the other miRNA, let-7a (Fig. S6A, *r*^2^ = 0.24), perhaps because of weaker repression signal imparted by the presence of endogenous let-7 in HeLa cells. Nonetheless, when plotting the combined results for all six miRNAs the performance was strong (Fig. 5D, *r*^2^ = 0.34).

**Fig. 6.**
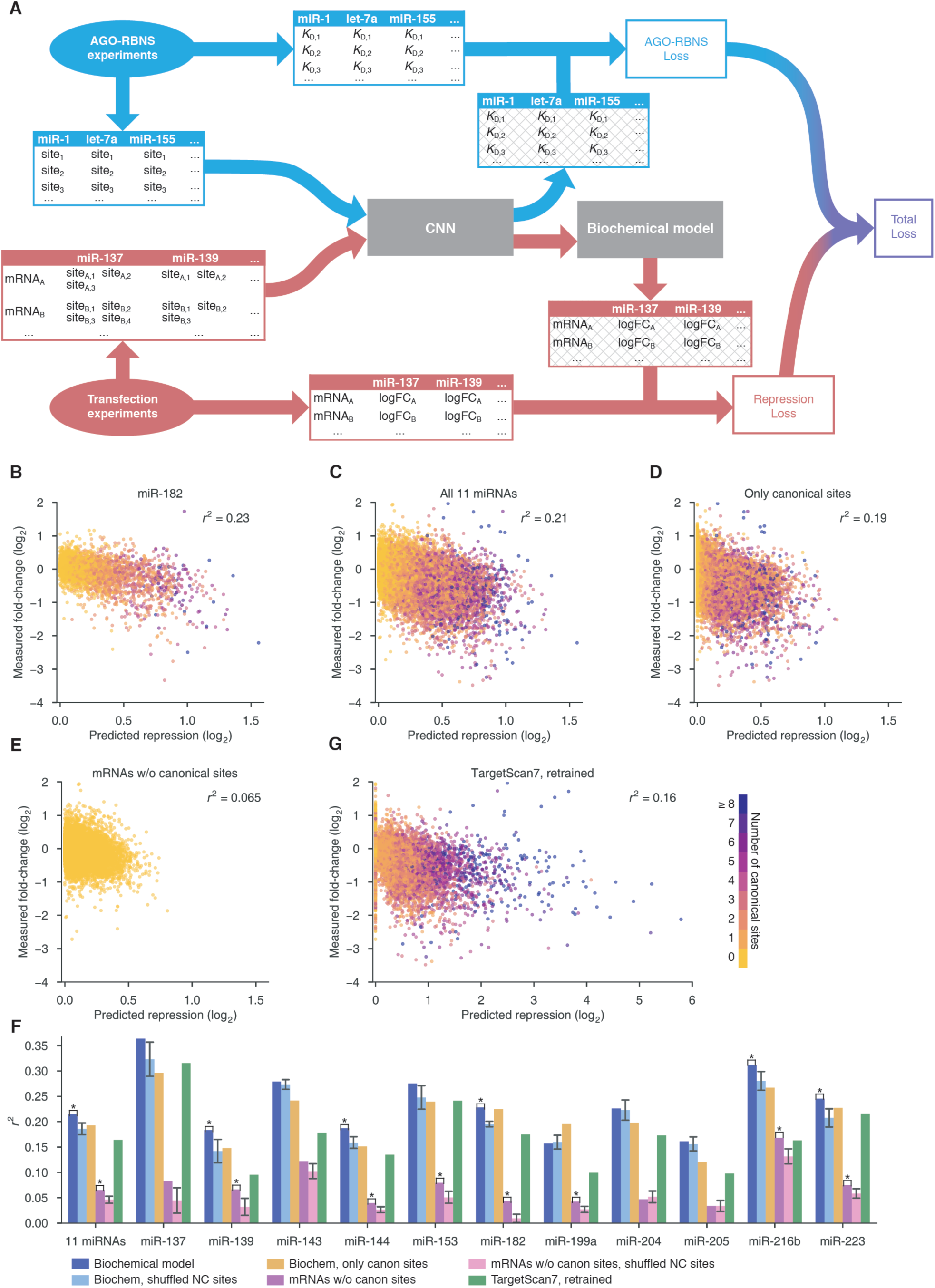
A CNN for predicting binding affinity from sequence. (**A**) Schematic of overall model architecture for training on RBNS data and transfection data simultaneously. “Loss” refers to squared loss. Tables with hash marks indicate model-predicted values, rather than experimentally measured values. (**B**) Typical test performance of the model when using the CNN-predicted relative *K*_D_ values. Results are shown for miR-182, chosen because among the 11 miRNAs tested, it had a median *r*^2^ value (Fig. S8A). Otherwise, as in Fig. 5C. (**C**) Test performances of the model when using the CNN-predicted relative *K*_D_ values for the 11 miRNAs without RBNS data. Otherwise, as in (B). (**D**) Performance of the model when refit using CNN-predicted relative *K*_D_ values for only 12-nt sequences with one of the six canonical site-types. Otherwise, as in (B). (**E**) Performance of the model when only considering mRNAs without one of the six canonical sites. Likelihood ratio test calculated as in Fig. 5F with *p* < 10^-15^. Otherwise, as in (B). (**F**) The contributions of cognate noncanonical sites to performance and comparison of performance to that of TargetScan. This panel is as in Fig. 5G, except it shows the results when using CNN-predicted relative *K*_D_ values for the 11 miRNAs without RBNS data (using 250 random shuffles rather than all possible permutations). (**G**) Test performance for the TargetScan model retrained on the 16 transfection datasets, showing combined results for the 11 miRNAs of (F). The *x*-axis was truncated at 6, which excluded 3 outlier points; otherwise, as in (B).

When provided *K*_D_ values for only the 12-nt *k*-mers that contained one of the six canonical sites, the model captured somewhat less variance (Fig. 5E, *r*^2^ = 0.32), and conversely when considering transcripts that lacked any canonical sites, the model had vastly diminished but statistically significant predictive power (Fig. 5F, *r*^2^ = 0.051, *p* < 10^-15^, likelihood ratio test). These results agreed with our previous analyses indicating that noncanonical sites can mediate intracellular repression (Fig. 3D–I). Alternatively, noncanonical sites might act as a proxy for a feature that correlates with repression. For example, noncanonical sites in higher AU context tend to have better *K*_D_ values (Fig. 4B, Fig. S5A–E) and could therefore be a reflection of UTR AU content. To control for this possibility, we repeated the analysis after replacing the noncanonical sites (and their *K*_D_ values) of each miRNA with those of another miRNA, performing this shuffling and reanalysis for all 265 possible shuffle permutations. When using each of these shuffled controls, performance decreased, both when considering all mRNAs and when considering only those without a canonical site to the cognate miRNA (Fig. 5G, five miRNAs), as expected if the modest improvement conferred by including noncanonical sites were due, at least in part, to miRNA pairing to those sites. This advantage of cognate over shuffled noncanonical sites was largely maintained when evaluating the results for individual miRNAs (Fig. 5G). Together, our results showed that noncanonical sites can mediate intracellular repression but that their impact is dwarfed by that of canonical sites because high-affinity noncanonical sites are not highly abundant within 3′ UTRs.

The performance of our biochemical model (*r*^2^ = 0.34) exceeded those of the 30 target-prediction algorithms (*r*^2^ ≤ 0.14) that were also tested on changes in mRNA levels observed in response to miRNA transfection (*13*). We reasoned that in addition to our biochemical framework and the use of experimentally measured affinity values, other aspects of our analysis might have contributed to this improvement. These included our evaluation of the model on all mRNAs, rather than only mRNAs with a 7–8-nt canonical 3′-UTR site, and the improved transfection datasets used to evaluate our model (which had stronger signal over background compared to microarray datasets used to train and test previous target-prediction algorithms). To evaluate the contributions of these differences on the improved performance of our model, we generated transfection datasets for 11 additional miRNAs and retrained the latest TargetScan model on the collection of 16 miRNA-transfection datasets (omitting the let-7a dataset to avoid the complications of the endogenously expressed let-7 miRNAs), putting aside one dataset each time in a 16-fold cross-validation. Training and testing TargetScan on improved datasets and considering all mRNAs did increase its measured performance, yielding an *r*^2^ of 0.24 for the five miRNAs with AGO-RBNS data, excluding let-7a (Fig. 5H), which exceeded the *r*^2^ of 0.14 previously reported for TargetScan. Nonetheless, the biochemical model still outperformed the retrained TargetScan by 15–55%, depending on the miRNA (Fig. 5G), which showed that use of measured affinity values in a biochemical framework substantially increased prediction performance.

### Convolutional neural network for predicting site *K*_D_ values from sequence

Our findings that binding preferences differ substantially between miRNAs and that these differences are not well predicted by existing models of RNA duplex stability in solution posed a major challenge for applying our biochemical framework to other miRNAs. Because performing AGO-RBNS for each of the known miRNAs would be impractical, we attempted to predict miRNA–target affinity from sequence using the relative *K*_D_ values and miRNA-transfection data already in hand. Bolstered by recent successful applications of deep learning to predict aspects of nucleic acid biology from sequence (*36*–*38*), we chose a convolutional neural network (CNN) for this task.

The overall model had two components. The first was a CNN that predicted *K*_D_ values from miRNA and 12-nt *k*-mers (Fig. S7A), and the second was the previously described biochemical model that links intracellular repression with *K*_D_ values (Fig. 6A). The training process simultaneously tuned the both the neural network weights and the parameters of the biochemical model to fit both the relative *K*_D_ values and the mRNA repression data, with the goal of building a CNN that accurately predicts the relative *K*_D_ values for all 12-nt *k*-mers of a miRNA of any sequence.

The input data for the CNN consisted of over 1.5 million *K*_D_ values from six AGO-RBNS experiments and 68,112 mRNA expression estimates derived from 4,257 transcripts in 16 miRNA transfection experiments (excluding let-7a to avoid the complications of the endogenously expressed let-7). Five miRNAs had data in both sets. Because some repression was attributable to the passenger strands of the transfected duplexes (Fig. S7B), the model considered both strands of each transfected duplex, which allowed the neural network to learn from another 16 AGO-loaded guide sequences. During training, we systematically left out the data for one of the 11 miRNA duplexes without AGO-RBNS data and trained on the remaining datasets, in an 11-fold cross-validation procedure. The predicted *K*_D_ values for each held-out dataset were then used as input for the biochemical model to predict repression for each of the 11 miRNAs (Fig. S8A, *r*^2^ = 0.16–0.36), with a median performance of *r*^2^ = 0.23 (Fig. 6B) and overall performance of *r*^2^ = 0.21 (Fig. 6C).

When the biochemical model was provided CNN-predicted *K*_D_ values for only the 12-nt *k*-mers that contained one of the six canonical sites, its performance decreased somewhat (Fig. 6, C and D, *r*^2^ = 0.21 and 0.19, respectively), and when considering transcripts that lacked any canonical sites, the model captured a minor but significant portion of the variance (Fig. 6E, *r*^2^ = 0.065, *p* < 10^-15^, likelihood ratio test). Likewise, an advantage was observed when using cognate over noncognate noncanonical sites, and these trends were maintained for most of the miRNAs when evaluated individually (Fig. 6G). These results mirrored those observed when using relative *K*_D_ values derived from AGO-RBNS, indicating that the CNN had some ability to identify effective noncanonical *K*_D_ sites for new miRNAs.

The biochemical model provided with CNN-predicted *K*_D_ values outperformed the retrained version of TargetScan, both when comparing the results for all 11 miRNAs in aggregate (Fig. 6C, F, *r*^2^ = 0.21 and 0.16, respectively) and when comparing them for each miRNA individually (Fig. 6G). Indeed, for miR-137 and miR-216b, the performance observed when using the *K*_D_ values predicted from the neural network rivaled that of the biochemical model applied to experimentally determined *K*_D_ values (Fig. 6G, Fig. 5G). In some cases, the ability to account for the effects of both strands of the miRNA duplex might offset the disadvantages of using less accurate *K*_D_ values.

### Insights into miRNA targeting

Our results provide new insight into both the canonical and noncanonical miRNA site types. For each miRNA, the canonical 8mer site was the highest-affinity site identified, illustrating its primacy in miRNA targeting. However, the canonical 7mer-m8 was the not always the second-most effective site; miR-155 had one noncanonical site with greater affinity than that of this canonical site, and miR-124 had five (Fig. 2, Fig. S2). Moreover, four of the six miRNAs had noncanonical sites with greater affinity than that of the canonical 7mer-A1 sites. Indeed, miR-124 had 29 noncanonical sites with greater affinity than that of the canonical 7mer-A1site and 40 noncanonical sites with greater affinity than that of the canonical 6mer site. (Fig. 2, Fig. S2).

The observation that canonical sites are not necessarily those with the highest affinity raises the question of how canonical sites are distinguished from noncanonical ones and whether making such a distinction is useful. Our results show that two criteria readily distinguished canonical sites from noncanonical ones. First, the six canonical site types were the only ones identified for all six miRNAs, whereas the noncanonical site types were typically identified for only one miRNA, and never for more than three. Second, the four highest-affinity canonical sites occupied most of the specifically bound AGO2, even for miR-124, which had the largest and highest-affinity repertoire of noncanonical sites (Fig. 1F, Fig. 2). This greater role for canonical sites was presumably because perfect pairing to the seed region is the most efficient way to bind the silencing complex; to achieve equivalent affinity, the noncanonical sites must be longer and are therefore less abundant. For example, although the miR-124 7mer-m8 site had lower affinity than a 11-nt noncanonical site, the canonical 7-nt site occupied much more AGO2–miR-124 because of its 256-fold greater abundance. The ubiquitous function and more efficient binding of canonical sites explains why these site types have the greatest signal in meta-analyses of site conservation, thereby explaining why they were the first site types to be identified (*30*) and justifying the continued distinction between canonical and noncanonical site types.

The potential role of pairing to miRNA nucleotides 9 and 10 has been controversial. Although some target-prediction algorithms (such as TargetScan) do not reward pairing to these nucleotides, most algorithms assume that such pairing enhances site affinity. At the other extreme, a biochemical study reports that such pairing can reduce site affinity (*6*). We found that pairing to nucleotides 9 and 10 neither enhanced nor diminished affinity in the context of seed matched sites (Fig. 4), whereas pairing to nucleotides 9 and 10 enhanced affinity in the context of 3′-only sites (Fig. 2C, D). These results support the idea that extensive pairing to the miRNA 3′ region unlocks productive pairing to nucleotides 9–12, which is otherwise inaccessible (*1*). Moreover, we found that although the nucleotides at target positions 9 and 10 seem unable to pair to the miRNA in the context of most canonical sites, nucleotide composition at positions 9 and 10 can have a dramatic influence on the affinity of canonical sites through an effect on site accessibility (Fig. 4).

The success of our biochemical model in predicting how each mRNA would respond to a transfected miRNA (Fig. 5) supports the conclusion that site-binding affinity is the major determinant of miRNA-mediated repression and that noncanonical sites measurably contribute to this repression in the cell. When developing our model, we also considered an alternative version in which the extent of repression scaled with the probability of binding at least one silencing complex, rather than with the average number of silencing complexes bound. However, the current model was more predictive for repression (Fig. S8B), which supported the idea that each additional bound AGO molecule contributes additional repression.

The biochemical parameters fit by our model provided additional insights into miRNA targeting. In the framework of our model, the fitted value of 1.01 ± 0.03 observed for the parameter *b* suggested that, in the absence of the UTR-length effect, a typical mRNA bound to an average of one silencing complex will experience a doubling of its decay rate, which would halve its abundance. In the concentration regimes of our transfection experiments, this occupancy can be achieved with two to three median 7mer-m8 sites. Another parameter, which was fit separately for each transfection experiment, was *a*_*t*_, i.e., the intracellular concentration of AGO2 loaded with the transfected miRNA and not bound to a target site. This parameter was expected to be affected, in part, by the abundance of target sites for that miRNA in the transcriptome, which is a known correlate of miRNA-targeting efficacy (*28, 39*). Indeed, we observed the expected relationship between the fitted *a*_*t*_ and predicted target-site abundance within the transcriptome (Fig. S8C).

Some other features known to correlate with targeting efficacy are also captured by our biochemical model. Indeed, the contribution of certain features, such as site type (*3*), predicted seed-pairing stability (*28*), and nucleotide identities at specific miRNA/site positions (*13*), are expected to be captured more accurately in the miRNA-specific *K*_D_ values of the 12-nt *k*-mers than when generalized across miRNAs. One feature that was not captured in our model is the evolutionary conservation of sites, although its utility is expected to decline as the biochemical model becomes more accurate. Another feature important for ∼5% of the sites is the contribution of supplementary pairing to the miRNA 3′ region (*3*). AGO-RBNS using libraries designed to query this additional pairing will provide the information needed to accurately incorporate it into our model. We suspect, however, that the most promising approach for improving our model will be to perform AGO-RBNS with additional miRNAs, as each additional site-affinity profile will facilitate direct modeling of the regulatory effects of that miRNA in vivo. Acquiring more site-affinity profiles for miRNAs with diverse sequences will also enable improvement of the CNN, whereby the complete RNA–RNA affinity landscape, as specified by AGO within this essential biological pathway, might ultimately be computationally reconstructed.

## ACKNOWLEDGEMENTS

We thank K. Heindl, T. Eisen, and T. Bepler for helpful discussions, and members of the Bartel lab for comments on this manuscript. This work was supported by NIH grants GM118135 (D.P.B.) and GM123719 (N.B.). D.P.B. is an investigator of the Howard Hughes Medical Institute. Sequencing data will deposited in Gene Expression Omnibus, and computational tools will be deposited in GitHub.

## SUPPLEMENTARY MATERIALS

Experimental Materials and Methods

Mathematical and Computational Methods

Figures S1 to S9

Tables S1 and S2

Data S1 and S2

